# Online interference of declarative memory on fast and slow adaptive processes in force field motor learning

**DOI:** 10.1101/2024.12.20.629631

**Authors:** J. L. Rudolph, L. P. J. Selen, W. P. Medendorp

## Abstract

Error-based motor adaptation is currently understood as a dual-rate process involving a fast adaptive process that learns quickly but also decays rapidly and a slow process that learns slowly but has good retention. While the fast process is typically categorized as procedural learning, recent evidence suggests that it relies on the declarative memory system. To test this hypothesis, we investigated in what manner a declarative memory task interferes with two processes that supposedly underly force field adaptation in reaching. This declarative memory task, which involved learning of a list of words, was assessed through either recognition or recall, and was compared to a non-declarative, vowel counting task, using a within-subject design (n=32). We employed a Bayesian hierarchical dual-rate process model to capture the observed force compensation across trials, expecting that the parameters of the fast process would be affected by the declarative memory task. We examined the 95% highest density interval of the posterior distribution of the difference between the experimental and control condition for each parameter. While most parameters remained unaffected by the declarative memory task, the retention rate of the fast process showed a hint of reduction, suggesting a complex interplay between declarative memory and ongoing motor adaptation processes.

## Introduction

Long-term memory is typically divided into declarative or explicit memory and non-declarative or implicit memory (Atkinson & Shiffrin, 1968; Squire, 2004). Explicit memory refers to conscious recollection of facts and events; implicit memory is automatic and includes knowledge of skills and how to perform tasks, also called procedural memory. Motor learning is seen as a form of procedural memory, defined as the process of (re)gaining or retaining a given level of motor performance.

Motor learning can be studied in the context of reach adaptation in which participants learn to compensate for a systematic perturbation, for example a visuomotor rotation (Cunningham, 1989; see Krakauer et al., 2019 for review) or a force applied to the hand (Shadmehr & Mussa-Ivaldi, 1994; see Krakauer et al., 2019 for review). Healthy people show a monotonic improvement in such tasks that is initially rapid but slows when approaching plateau performance. This behavioral learning curve is well described by an exponential curve, suggesting that the amount of improvement on each trial is proportional to the error (Shadmehr et al., 2010).

Smith et al., (2006) suggested that reduction of error engages two independent adaptive processes, with different dynamics. Their suggestion is based on a paradigm that evokes *spontaneous recovery*, the observation that an initially learned behavior re-emerges after it has been extinct for a while (Pavlov, 1927; Rescorla, 2004). Smith and colleagues performed a force field adaptation experiment with reaching movements. In their paradigm, they recorded the participant’s forces while they first learned to compensate for one force field for an extensive period, followed by a brief exposure to the opposite force. After this brief exposure, compensation for the first force field seems to have disappeared. In the next phase, participants made clamped reaching movements, without a perturbing force field, thereby removing any error feedback. In this phase, compensatory forces gradually changed from aligning with the second force field to aligning with the first force field, i.e., a recovery of the compensation for the first force field. This gradual recovery of compensation was predicted by a dual-rate model with a fast process that learns quickly, accounting for early adaptation, but also forgets rapidly and a slow process that learns slowly and forgets slowly.

Because the fast adaptive process is mainly driven by large errors, Keisler and Shadmehr (2010) suggested that it shares resources with the declarative memory system. The idea is that large errors result in awareness of the perturbation (Malfait & Ostry, 2004), which, driven by such explicit processing, engages the declarative memory system. In support, Keisler and Shadmehr (2010) found that imposing a declarative memory task (word-list learning task) after completion of a motor adaptation task had a negative effect on the motor memory associated with the fast adaptation process, but not with the slow adaptation process. Hence, a word-learning task can cause retrograde interference on a motor memory.

Taylor et al. (2014) also suggested that the fast adaptation process relies on declarative memory. These authors asked participants to report the aiming direction of their intended movement in a visuomotor adaptation experiment. The reports, which provide a measure of an explicit strategy, were found to follow the dynamics of the fast adaptation process (McDougle et al., 2015). Furthermore, McDougle et al. (2022) described alterations of the fast adaptive process in a case report of a patient with near-complete bilateral loss of her hippocampi, known to be involved in declarative memory. Recently, Sutter et al. (2024) found that early-stage Alzheimer’s disease, which is characterized by an initial decline in declarative memory, affects the fast but not the slow adaptive processes in motor learning. Hence, all this work has led to the notion that motor adaptation not only recruits non-declarative, i.e. implicit, but also declarative memory processes.

Declarative memory processes are known to be impaired by both retrograde (or retroactive) and anterograde (or proactive) interference manipulations (Crowder, 1976; Gluck et al., 2014; Postman & Underwood, 1973). In turn, to establish a connection of the fast process in motor learning with declarative memory processes, requires a demonstration of both retrograde and anterograde interference effects. While Keisler & Shadmehr (2010) reported retrograde interference on the fast process, anterograde interference effects have not been examined. Therefore, in the current study we investigate how a declarative memory task interferes antero- and retrogradely with trial-by-trial force field reach adaptation in a spontaneous recovery paradigm.

## Methods

### Participants

For this study 33 participants were recruited via the university’s participation database system (SONA system). Each participant performed two sessions, which were less than 16 days apart (on average 4 days). One participant did not show up for the second session and was therefore replaced. The remaining 32 native Dutch, non-dyslexic participants (22 female, 10 male) were on average 25 years old (ranging from 17 to 66), had normal or corrected to normal vision, were right-handed, and had no motor impairments. The study used a within-subject design. Each participant was randomly assigned to one of eight groups, which were defined based on which declarative memory task was tested (recognition or recall, see below), whether the control task was performed before or after the memory task (i.e., in session 1 or 2) and the force field order tested in the memory and control task (CW-CCW or CCW-CW, respectively).

Further details on these groups are below. The duration of a session varied between 22 and 36 minutes. Participants were reimbursed with either a 15€ gift-card, or 1.5 participation points as part of the bachelor’s course programme (for students).

### Setup

Participants sat in front of a robotic manipulandum (3BOT) that was configured to move in the transversal plane (configured as a 2D-vBOT, Howard et al., 2009). Holding the handle at the endpoint of the manipulandum, participants made right-handed reaching movements while their forearm was supported by an air-sled to minimize friction. The robot recorded handle positions and generated forces at 1000 Hz.

To present visual stimuli into the participant’s workspace, participants viewed an LCD display via a mirror above the hand, which presented the start (grey circle, 1 cm radius), the target (yellow circle, 1cm radius), a cursor (red circle, 0.5 cm radius) representing current hand position as well as written reach performance feedback, all against a black background. Participants could not see their hand. The start location was presented at the center of the screen approximately 30 cm away from the participant’s body, on the body midline. The target location was presented 12 cm further away, also on the body midline. Performance feedback was presented between trials, 2 cm below the target location.

### Reach task

Participants made forward reaching movements from the start location to the target location. In each trial, participants were required to hold the hand cursor stationary (speed < 2.5 cm/s) for 100 ms within the start location, before the target appeared instructing the participants to initiate the reach. Participants were instructed to move through the target. If participants did not initiate the movement within 600 ms after target presentation, a feedback message stating “react faster” appeared on the screen and the trial was repeated. If participants stopped the reach (speed < 2.5 cm/s) without traversing a displacement of 12 cm (i.e., radial distance from start to target), the trial was aborted and repeated, with the feedback “smoothness” presented. A reach was considered successful if the participant moved through the target within 250–350 ms after movement onset (i.e., when speed > 2.5 cm/s). In this case, the feedback message “correct” appeared on the screen and the visual target exploded into a firework of 20 circles of two randomly chosen colors that slowly decreased in size and moved radially away from the target location, accompanied by a pleasant sound. If participants did not reach through the target within the required time interval, the message ‘Move Faster’/’Move Slower’ appeared, while the target jittered at its location (red for ‘Move faster’ and blue for ‘Move slower’), accompanied by the same pleasant sound (indicating that the target was still hit). When participants missed the target, the message “Accuracy” was shown, together with a less pleasant sound. Furthermore, the target width increased such that it includes the location where the cursor had crossed the 12 cm bound, before shrinking back to its original size. When the handle speed dropped below 2.5 cm/s, the robot guided the hand back to the start location in 0.7s following a bell-shaped velocity profile.

The experiment comprised three types of reach trials: *null trials, force field trials* and *error-clamp trials*. In null trials, the robot produced no forces. In force field trials, the robot produced a velocity-dependent curl force field. The force (*F*) depended on the 2D hand velocity (*ẋ, ẏ*), and a field constant 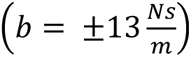, creating a force that is proportional to hand speed and perpendicular to the instantaneous movement direction:

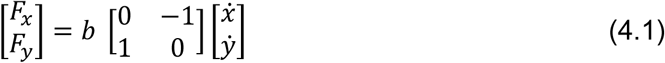

In error-clamp trials, the active compensation (i.e., adaptation) by the participant was measured by constraining the reaching movement to a straight path from the start position of the movement (hand speed > 2.5 cm/s) to the end of the movement (hand speed < 2.5 cm/s, Scheidt et al. 2000), with a stiffness of 3000 N/m and a damping constant of 25 Ns/m. Note, these error-clamps run through the center of the target location.

### Interference tasks

To test whether a declarative memory task interferes with force field adaptation, we used the “15 words test” (in Dutch: “15 woorden test”, Kalverboer & Deelman 1986, Version A). This declarative memory task contains 15 Dutch words which was adapted for our purposes to a recall and a recognition task with the aim to tax declarative memory differently (de Vanssay-Maigne et al., 2011; Scalici et al., 2017). The 15 words were first, one-by-one, shown on the screen, interleaved by three reach trials (see Figure 1). Subsequently, after every three reach trials, participants performing the recall task were asked to name one of the words (Figure1A, left panel). If the participant indicated they could not recall any more words, or had successfully recalled 15 words, the full word list was presented again, following the procedure above. This cycle of presenting and recalling was repeated 5 times. During the last round the participant was only asked to memorize 6 words due to paradigm constraints (see section Experimental Paradigm). Participants performing the recognition task were asked which words they recognized among a list of new words (see Figure 1A, middle panel). The 30 words (15 shown and 15 new words) from the recognition task (Kalverboer & Deelman 1986) were split into 5 sets of 6 words. This cycle of presenting and recognizing was also repeated 5 times.

As a control for the intervening declarative recall and recognition tasks, we used a vowel counting task, which was performed after every three reach trials (see Figure 1A, right panel). In this task participants viewed a string of letters on the screen and had to verbally report the number of vowels. The string length varied between 3 and 5 letters and could contain 0-3 vowels; we used 105 different strings.

### Experimental paradigm

Eight groups of 4 participants each engaged in the force field reaching task following a spontaneous recovery paradigm that embedded either the recall, the recognition or the vowel counting task (see Figure 1B). Two groups performed the recall task during the first session, and the vowel counting task during the second session. The groups differ in the order of force fields (CW or CCW first) in the spontaneous recovery paradigm. Two other groups performed the vowel counting task first. The same holds for the remaining four groups except they performed the recognition, not the recall task. Due to this counterbalancing, data can be aggregated in two groups: a recognition group (n=16, assessing the effect of intervening word recognition on force field adaptation versus control) and a recall group (n=16, the effect of word recall on the motor adaptation).

The reach paradigm with the vowel counting task contained 622 trials: 150 baseline trials, 252 trials for the first force field (either counterclockwise (CCW) or clockwise (CW)), 20 trials for the second opposite force field, 50 error-clamp trials to test for spontaneous recovery, and 150 washout trials (Figure 1B). The paradigm including the recall task was identical except that the number of trials during the first force field varied across participants, depending on the number of recalled words (Figure 1B blue, variable number of trials indicated by smaller dots). As a result, the total number of trials ranged from 680 to 731 trials (M = 710 trials). The paradigm including the recognition task had the same number of trials as with the vowel counting task.

In all three paradigms, every 8^th^ trial in the baseline, first and second force field phase, and washout phase was an error-clamp trial. Two additional error-clamp trials were interspersed after the second trial of the first and second force field to assess the level of early adaptation more thoroughly. From trial 108 onwards, every third trial was preceded by the presentation, recall, or recognition of a word (memory tasks), or a string of letters (vowel counting task) until the end of the second force field (see Figure 1B, interference task). The schedule of the interference task aligned the first and last recall and recognition phase of the words with the introduction of the first and second force field, respectively.

### Behavioral analysis

Offline data analyses were performed in MATLAB (2019b) and R (R Core Team, 2022). Position and force data was filtered at 12 Hz using a 5th order, zero lag, Butterworth filter. Filtered position data was differentiated to calculate velocity and speed. From the filtered data (criteria described above) movement onset was defined as the first timepoint where the hand speed exceeded 15% of the maximum speed during the outgoing movement, whereas movement end was defined as the first point where the speed dropped below 15% of maximum speed.

Two measures were used to quantify adaptation: the hand path error (HPE) and the adaptation index (AI). The HPE was calculated on trials without an error-clamp and was defined as the angle between the line connecting the hand’s starting position and the target position and the line connecting the hand’s start position and the hand’s position at maximum speed. For visualization purposes, the individual participant’s HPE of the CW-CCW group was sign-reversed to match with the CCW-CW group.

The AI quantifies the active compensation for the expected force field (Smith et al., 2006). It is computed by regressing, without an offset, the expected force estimated by Eq. (1) with the force exerted into the error-clamp by the participant. This provides a regression coefficient between -1 and 1. The sign was introduced to indicate the direction of compensatory force: positive CW, negative CCW. AIs were baseline corrected by subtracting the average AI obtained during the baseline phase of the experiment. Again, for visualization purposes, AIs were collapsed over CW and CCW force fields after flipping the sign of the CCW force field.

### Hierarchical Bayesian dual-rate model

To examine how the declarative memory task interfered with force field adaptation, we implemented a hierarchical version of the dual-rate model (Smith et al., 2006). This hierarchical model was inspired by the dual-rate model of Hulst et al. (2020) for visuomotor adaptation and Sutter et al. (2024) for force field learning. This model makes use of partial pooling in which the estimate of the parameters of individual participants are influenced by the estimate of other participants through the estimate of hyperparameters. In our model (for a schematic representation, see Figure 2), the observed adaptation indices (AI) of each participant (*n*), session (*s*) and trial (*t*) are assumed to be drawn from a normal distribution:

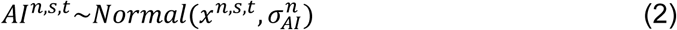

**Figure 2.**
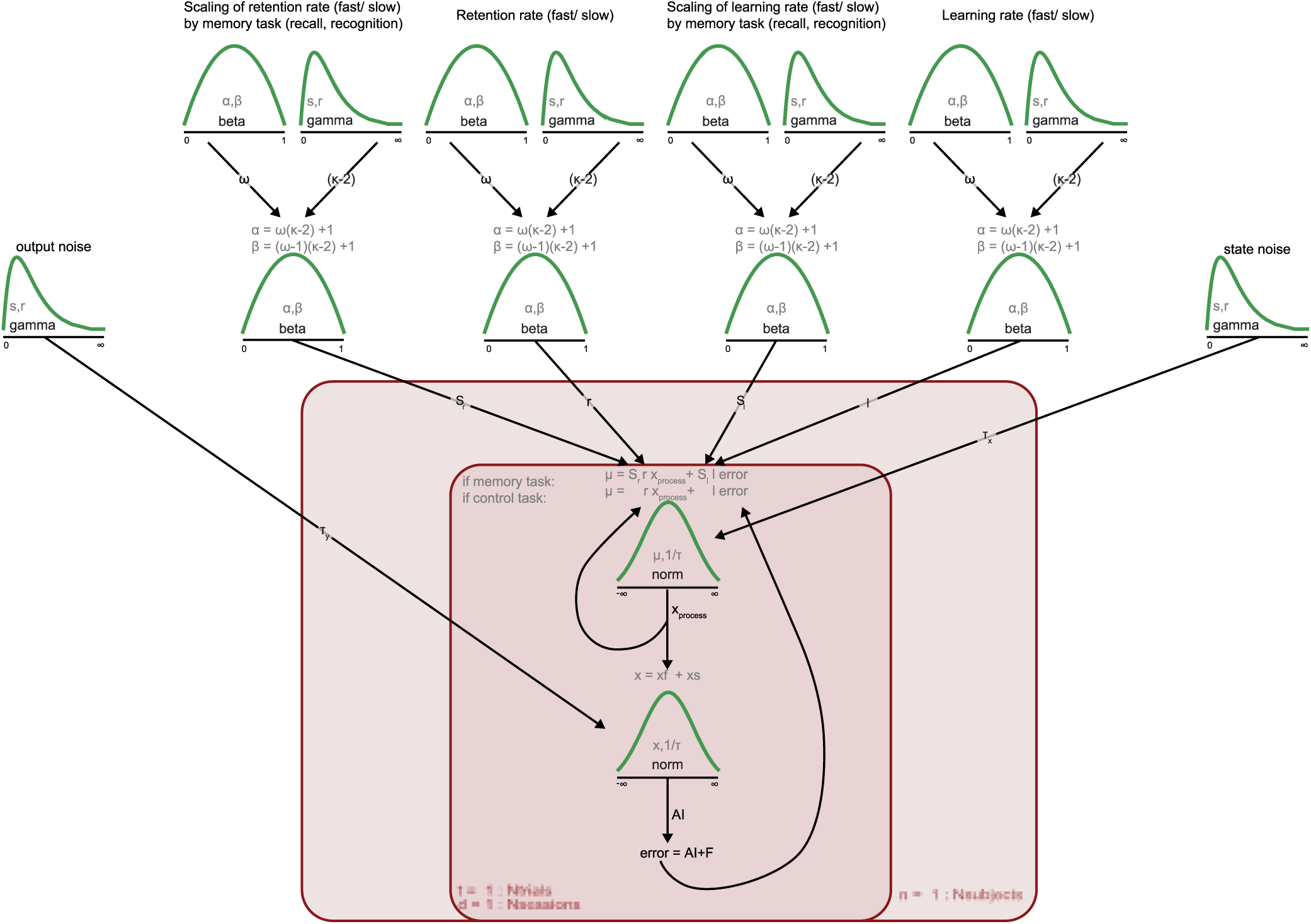
Scheme of the hierarchical Bayesian dual-rate model. parameters within a red square are drawn or calculated per participant/trial and session. The beta distributions represent the hyper-distributions from which the parameter values of individual participant distributions are drawn.

with a mean *x^n,s,t^* and a standard deviation 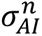. The precision of the observations 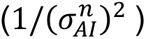 is drawn from a gamma distribution:

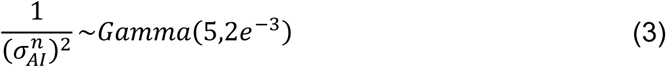

The state *x^n,s,t^* indicates the force a participant was planning to use on a certain trial and session, without the noise that occurred during the execution of the movement. This state is based on the sum of the two underlying processes:

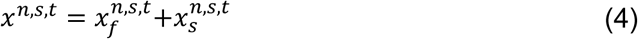

in which 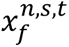 is the state of the fast process and 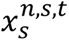 the state of the slow process, both drawn from a normal distribution:

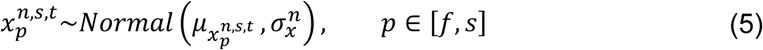

with a mean of 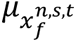 for the fast process and 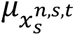 for the slow process, and their standard deviation 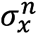, representing the process noise.

We decided to use the same process noise values for the fast and slow process to limit the number of parameters, just like Hulst et al. (2020) (also see Albert & Shadmehr (2018) for a comparison). The precision of the state noise 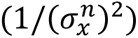 is related to the planning of the movement and is drawn from a gamma distribution:

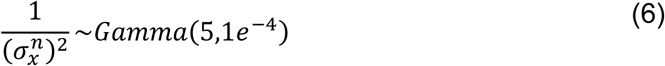

Both processes learn from error, which is defined as the sum of the compensatory force (i.e., *AI^n,s,t^*) and the (normalized) perturbing force (i.e., *F^n,s,t^*):

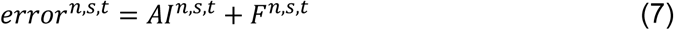

In the control condition, i.e. with an intervening vowel count task, the update of the fast and slow process depends on process-specific learning 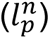 and retention 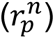 rates and follows the traditional model:

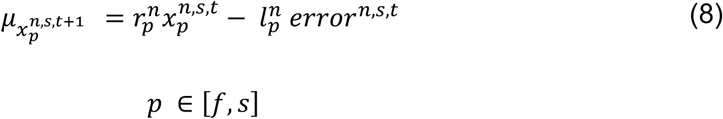

this 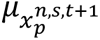 will be the mean of the fast and slow state of the next trial (Eq. 5). The parameter values of the learning and retention of the fast and slow process (here indicated as *q^n^*) are drawn, per participant, from population beta distributions:

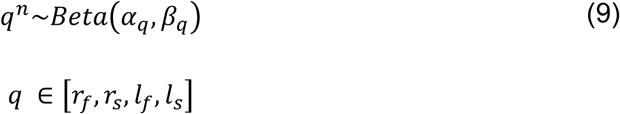

in which the parameters *α_q_* and *β_q_* of the beta distribution are calculated from the mode (*ω_q_* = 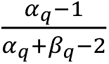) and concentration (*κ_q_* = *α + β*; Kruschke, 2015). The mode of the distribution (*ω_q_*) is drawn from a beta distribution (with parameters *a_q_* and *b_q_*) and the concentration of the distribution (*κ_q_* − 2) is drawn from a gamma distribution (with parameters *shape_q_* and *rate_q_*). These higher-order distributions assume that the estimates of learning and retention distributions of individual participants are drawn from population distributions (i.e., the hyper-distributions).

The priors on the mode (*ω_q_*) of the distribution and the concentration (*κ_q_* − 2) of the distribution were informative and based on the reported means and variances in Trewartha et al. (2014). We assume the mode of the learning and retention of the fast process to be centred around 0.5, and those of the slow process to be biased towards one of the extremes of the distribution (closer to 0 for the learning rate, and closer to one for the retention rate). Values of *a_q_* were therefore set to 2, 2, 2, and 0.7 (*r_f_,r_s_,l_f_,l_s_*), and values for *b_q_* were set to 2, 0.7, 2, and 2. We assume higher concentrations in the learning and retention of the slow process and lower concentrations in the learning and retention of the fast process. The values of the priors of the concentration (*κ_q_* − 2) were therefore set to 0.9, 2.5, 0.9, 2.5 for the *shape_q_* and 0.01, 0.02, 0.01, 0.02 for the *rate_q_*.

For the sessions that involved memory tasks (recall and recognition), we allowed the learning and retention rates of the fast and slow process to be only reduced relative to their values in the control task:

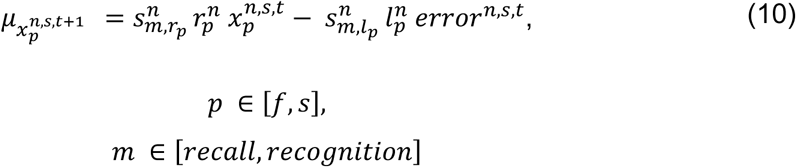

where 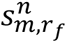 is the scaling of the fast retention rate, 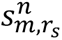 is the scaling of the slow retention rate, 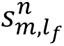 is the scaling of the fast learning rate, 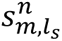 is the scaling of the slow learning rate, in which *N* represents the type of memory task. These scaling parameters were drawn from different beta distributions, one for each memory task per parameter:

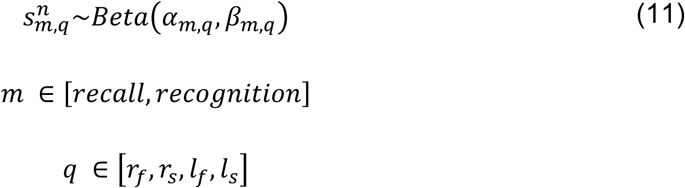

The modes of the scaling parameters were drawn from a uniform beta distribution (*ω_m,q_* ∼*Beta*(1,1)). The concentration (*κ_m_,_q_* − 2) of the scaling distributions were drawn from gamma distributions with the priors for the shape (*s_m,q_*) set to 2.5 and the priors for the rate (*r_m,q_*) set to 0.02 for all eight scaling parameters.

As described in previous work (Albert & Shadmehr, 2018), participants did not experience any error in error-clamp trials since their movement is restrained to a straight line from the start of the movement to the target. This was implemented into the model by setting the error to zero in Eqs. (8) and (10).

### Model fitting

For statistical analysis we used Markov Chain Monte Carlo (MCMC) sampling in R using Just Another Gibbs Sampler (JAGS; Plummer, 2016), to retrieve posterior distributions of parameter values. We ran the model with 4 chains letting it adapt for 30000 samples, followed by 120000 burn-in samples and then ran another 156250 samples to build the posterior distributions. Every 15^th^ sample was saved to reduce storage size and autocorrelation between samples (i.e., thinning).

Initial values (see Table 1, grey values in brackets) for the learning and retention rates of the hyperparameters were drawn from uniform distributions of the 95% HDI of the posterior distributions that were sampled from a pilot data set (n = 16) with a similar experimental paradigm to decrease the required number of samples to reach convergence. For every parameter, we first visually inspected convergence based on the diagnostic plots. Then, we consulted the effective sample size (ESS), the potential scale reduction factor (PSRF) and the Monte Carlo standard error (MCSE; see Table 2).

**Table 1.**
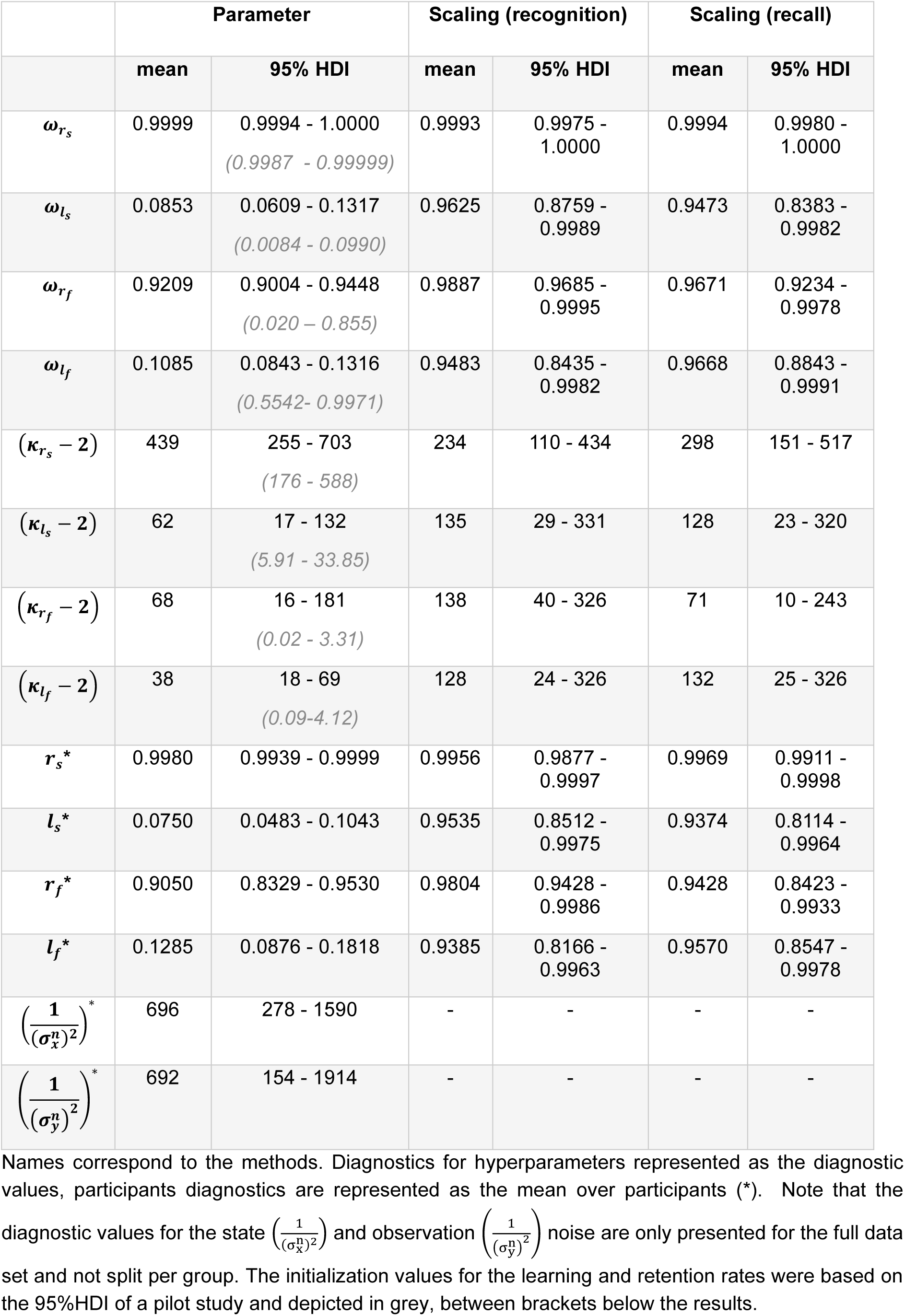
Mean and 95% HDI of the relevant parameters.

|  | Parameter |  | Scaling (recognition) |  | Scaling (recall) |  |
| --- | --- | --- | --- | --- | --- | --- |
|  | mean | 95% HDI | mean | 95% HDI | mean | 95% HDI |
| $\omega_{r_s}$ | 0.9999 | 0.9994 - 1.0000<br>(0.9987 - 0.99999) | 0.9993 | 0.9975 - 1.0000 | 0.9994 | 0.9980 - 1.0000 |
| $\omega_{l_s}$ | 0.0853 | 0.0609 - 0.1317<br>(0.0084 - 0.0990) | 0.9625 | 0.8759 - 0.9989 | 0.9473 | 0.8383 - 0.9982 |
| $\omega_{r_f}$ | 0.9209 | 0.9004 - 0.9448<br>(0.020 - 0.855) | 0.9887 | 0.9685 - 0.9995 | 0.9671 | 0.9234 - 0.9978 |
| $\omega_{l_f}$ | 0.1085 | 0.0843 - 0.1316<br>(0.5542- 0.9971) | 0.9483 | 0.8435 - 0.9982 | 0.9668 | 0.8843 - 0.9991 |
| $(\kappa_{r_s} - 2)$ | 439 | 255 - 703<br>(176 - 588) | 234 | 110 - 434 | 298 | 151 - 517 |
| $(\kappa_{l_s} - 2)$ | 62 | 17 - 132<br>(5.91 - 33.85) | 135 | 29 - 331 | 128 | 23 - 320 |
| $(\kappa_{r_f} - 2)$ | 68 | 16 - 181<br>(0.02 - 3.31) | 138 | 40 - 326 | 71 | 10 - 243 |
| $(\kappa_{l_f} - 2)$ | 38 | 18 - 69<br>(0.09-4.12) | 128 | 24 - 326 | 132 | 25 - 326 |
| $r_s^*$ | 0.9980 | 0.9939 - 0.9999 | 0.9956 | 0.9877 - 0.9997 | 0.9969 | 0.9911 - 0.9998 |
| $l_s^*$ | 0.0750 | 0.0483 - 0.1043 | 0.9535 | 0.8512 - 0.9975 | 0.9374 | 0.8114 - 0.9964 |
| $r_f^*$ | 0.9050 | 0.8329 - 0.9530 | 0.9804 | 0.9428 - 0.9986 | 0.9428 | 0.8423 - 0.9933 |
| $l_f^*$ | 0.1285 | 0.0876 - 0.1818 | 0.9385 | 0.8166 - 0.9963 | 0.9570 | 0.8547 - 0.9978 |
| $\left(\frac{1}{(\sigma_x^n)^2}\right)^*$ | 696 | 278 - 1590 | - | - | - | - |
| $\left(\frac{1}{(\sigma_y^n)^2}\right)^*$ | 692 | 154 - 1914 | - | - | - | - |

Names correspond to the methods. Diagnostics for hyperparameters represented as the diagnostic values, participants diagnostics are represented as the mean over participants (\*). Note that the diagnostic values for the state $\left(\frac{1}{(\sigma_x^n)^2}\right)$ and observation $\left(\frac{1}{(\sigma_y^n)^2}\right)$ noise are only presented for the full data set and not split per group. The initialization values for the learning and retention rates were based on the 95%HDI of a pilot study and depicted in grey, between brackets below the results.

**Table 2.** MCMC diagnostics for the relevant parameters.

|  | Parameter |  |  | Scaling (recognition) |  |  | Scaling (recall) |  |  |
| --- | --- | --- | --- | --- | --- | --- | --- | --- | --- |
|  | ESS | PSRF | MCSE | ESS | PSRF | MCSE | ESS | PSRF | MCSE |
| $\omega_{r_s}$ | 40818.9 | 1.00 | <0.0001 | 16787.9 | 1.00 | <0.0001 | 28121.4 | 1.00 | <0.0001 |
| $\omega_{l_s}$ | 1483.0 | 1.07 | 0.0006 | 1625.8 | 1.01 | 0.0008 | 1394.1 | 1.03 | 0.0012 |
| $\omega_{r_f}$ | 472.4 | 1.04 | 0.0005 | 657.4 | 1.01 | 0.0003 | 521.8 | 1.02 | 0.0009 |
| $\omega_{l_f}$ | 2652.2 | 1.01 | 0.0002 | 1034.4 | 1.00 | 0.0013 | 1431.4 | 1.00 | 0.0008 |
| $(\kappa_{r_s} - 2)$ | 2998.2 | 1.02 | 2.1000 | 2513.4 | 1.01 | 1.6700 | 14037.9 | 1.00 | 0.7960 |
| $(\kappa_{l_s} - 2)$ | 1084.1 | 1.06 | 0.9040 | 24482.9 | 1.00 | 0.5080 | 23369.6 | 1.00 | 0.5140 |
| $(\kappa_{r_f} - 2)$ | 256.2 | 1.10 | 2.7500 | 3156.1 | 1.00 | 1.3400 | 831.9 | 1.01 | 2.2100 |
| $(\kappa_{l_f} - 2)$ | 1355.5 | 1.06 | 0.3600 | 21420.4 | 1.00 | 0.5420 | 16013.2 | 1.00 | 0.6250 |
| $r_s^*$ | 11627.8 | 1.01 | <0.0001 | 7484.1 | 1.01 | 0.0001 | 9374.8 | 1.01 | <0.0001 |
| $l_s^*$ | 841.0 | 1.04 | 0.0006 | 2126.6 | 1.01 | 0.0009 | 1835.4 | 1.02 | 0.0012 |
| $r_f^*$ | 378.9 | 1.05 | 0.0017 | 1155.5 | 1.01 | 0.0005 | 748.8 | 1.04 | 0.0017 |
| $l_f^*$ | 813.3 | 1.04 | 0.0010 | 1346.9 | 1.00 | 0.0013 | 1743.8 | 1.00 | 0.0010 |
| $\left(\frac{1}{(\sigma_x^n)^2}\right)^*$ | 421.1 | 1.08 | 24.6272 | - | - | - | - | - | - |
| $\left(\frac{1}{(\sigma_y^n)^2}\right)^*$ | 878.2 | 1.06 | 20.0022 | - | - | - | - | - | - |

Names correspond to the methods. Diagnostics for hyperparameters represented as the diagnostic values, participants diagnostics are represented as the mean over participants (\*). Note that the diagnostic values for
the state $\left(\frac{1}{(\sigma_x^n)^2}\right)$ and observation $\left(\frac{1}{(\sigma_y^n)^2}\right)$ noise are only presented for the full data set and not split per group.
ESS: estimated Sample Size; PSRF: Potential Scale Reduction Factor; MCSE: Monte Carlo Standard Error

To infer whether the scaling of the learning and retention rates for the declarative memory tasks relative to the vowel counting task were of significance, we simulated participant behavior based on the experimental paradigm using the modes of the participant specific parameter distributions but kept the scaling parameters at one. Next, we used the same MCMC sampling method on this simulated dataset, creating a posterior distribution over the parameters, including the scaling parameter, knowing that there is no effect of the memory task. We then compared posterior scaling parameter distributions based on the experimental data and based on the simulations, using a Region of Practical Equivalence (ROPE) analysis (described in Kruschke 2015). We defined our ROPE based on the upper bound of the 90% HDI of the posterior distribution of the simulated data and set the boundaries to contain 10% of the samples of this posterior distribution. So, the bounds were set to the point containing 85% of the samples to the point containing 95% of the samples of the posterior distributions of the simulated data (see Figure 6, dashed lines).

To test how the posterior parameter distributions of the data accounted for the participant data, we performed a posterior predictive check. We performed forward simulations of the dual-rate model with randomly drawn parameters (n = 5000) from the posterior distributions, including the noise terms.

## Results

Participants performed a declarative memory task (learning a word-list, testing based on recall or recognition) during force field adaptation following a spontaneous recovery paradigm. As a control, participants performed the same reach adaptation task but with a vowel counting task instead of the word recall/recognition. The paradigm entailed a baseline phase, a long phase of one force field (CCW or CW), followed by a short phase of the opposite field, an error-clamp phase, and a washout phase. In five repeating cycles, the word list-task intervened every third trial with the motor adaptation task. Participants performed well on the word recognition task, with scores ranging between 80% and 100% correct (*M* = 96.0 %, *SD* = 6.3%). Performance in the word recall task was lower, with scores improving from 56% in the first cycle to 90% correct in the fourth cycle. In the control vowel counting task participants made virtually no errors, with scores between 95% and 100%.

To investigate if these tasks interfere with the motor adaptation process, we computed the hand path error (HPE, see Methods) to assess the trial-by-trial changes of the reach trajectories. Figure 3A and D illustrates the HPE (mean ± SEM) for the recognition and recall groups, in combination with the control (vowel counting), with each curve color-coded by the interfering task. The HPE straddles around zero during the baseline phase. With the onset of the first force field (trial 151), there is a large excursion away from baseline, which returns to baseline in the subsequent trials, following an exponential decay. With the sign switch of the force field (trial 403 in Fig 3C, and 528 in Fig 3D), the HPE again shows a large error, in the opposite direction, with a magnitude about twice the size as during the onset of the first force field, caused by the combined aftereffect and the opposite force of the new perturbation. Again, the HPE returns to baseline as learning proceeds. The HPE was not assessed during the subsequent error-clamp phase and was annulled during the final wash-out phase.

**Figure 3.**
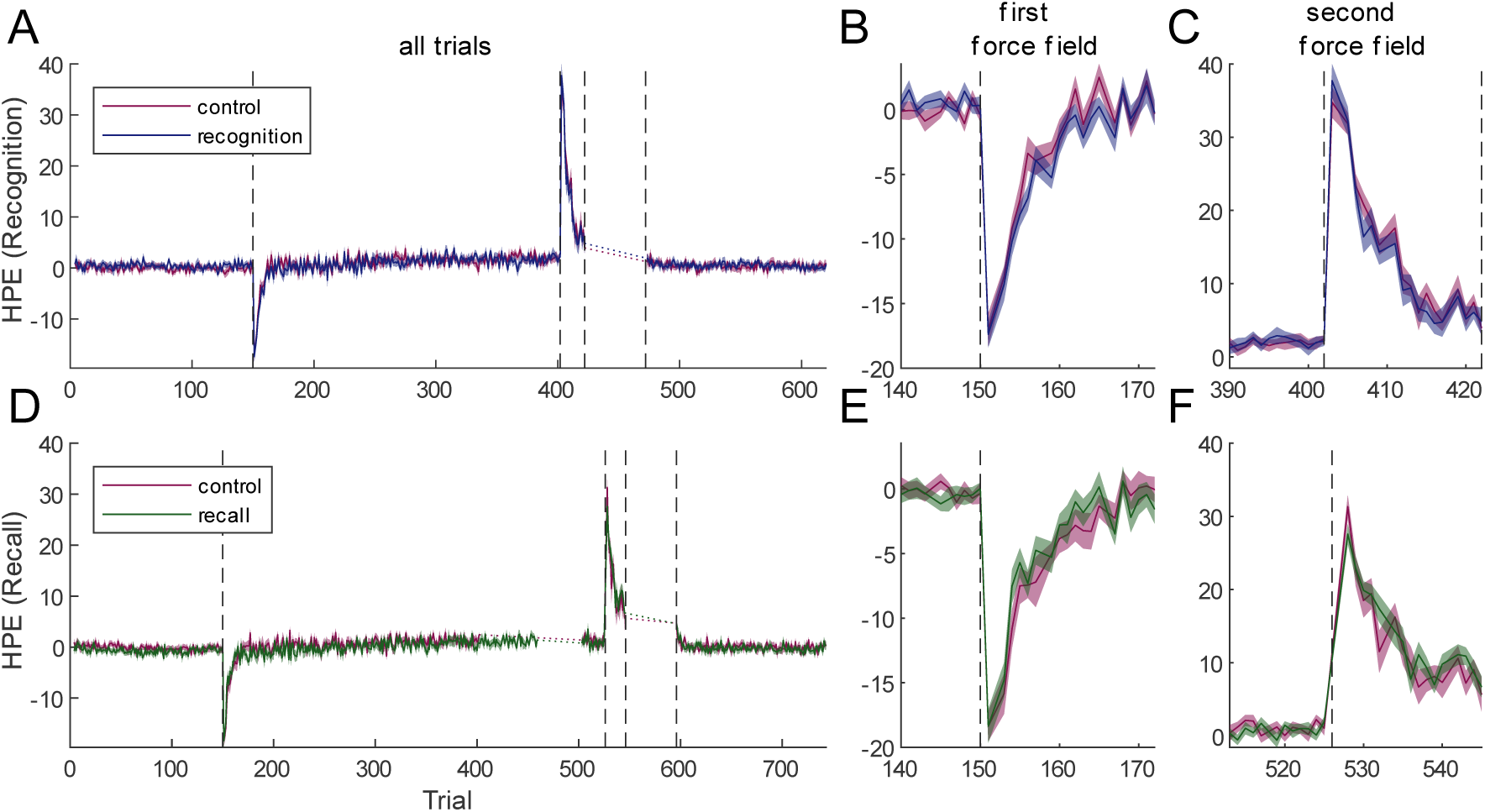
Hand path error (in degrees) across trials for the memory (green and blue) and control (purple) tasks for the recognition group (**A-C.**) and the recall group (**D-F.**). **A,D**. HPE for all trials. **B, F**. HPE for the trials around the introduction of the first force field, **C, E**. HPE for the trials around the switch between the first and second force field.

Neither panel A nor D in Fig 3 shows striking differences between the learning curves associated with the control and the respective memory task. Recall that the first declarative memory tests were aligned to the introduction of the first force field. If the fast process is impaired by these declarative memory tasks, one would expect the learning process to be slower compared to the control. Fig 3B and E, which zoom in on these differences, do not demonstrate any consistent effect. In support, post-hoc two-sided paired t-tests did not yield a significant difference between the HPEs in the control and memory session from trial 140 to trial 172 for both the recall and the recognition group (p>0.056, Bonferroni-corrected). A similar observation can be made for the HPE in response to the onset of the second force field (Fig 3C and F; p>0.059, Bonferroni-corrected). Finally, if the slow process was affected by the declarative memory task, one would expect differences in HPE to appear at the asymptote of the first force field, which was not visually present (Figure 3A for recognition and D for recall).

The observations in Figure 3 are corroborated in Figure 4, showing the changes in the adaptation index (AI ± SEM) – a measure of the compensatory force into the error-clamps – over the course of the experiment. In both groups, participants learn to compensate for the force fields, showing nearly full compensation (i.e., AI of ∼1) for the first force field after about 20 trials (see Figure 4A and E), irrespective of the intervening declarative memory task. The picture differs after the switch to the second force field, during which the Recognition group still reaches levels of full compensation, while the Recall group compensates by about half. As a result, during the spontaneous recovery phase, the AI rebounds (by about a change of 0.8), leading to higher levels of AI in the Recall than the Recognition group (Figure 4D and H). When visually inspecting the learning curves of the control and memory tasks, in neither the recall nor the recognition group there are obvious differences between the two learning curves. There is no reason to expect differences in the control task between the recall and recognition group, therefore we can only infer within-participant effects (see the hierarchical model results). During the spontaneous recovery phase, close scrutiny suggests that the curve associated with the declarative memory task lies slightly above the control curve, both for the recognition and recall group. This could indicate a lower retention of the second force field by the fast process due to the intervening declarative memory task.

**Figure 4.**
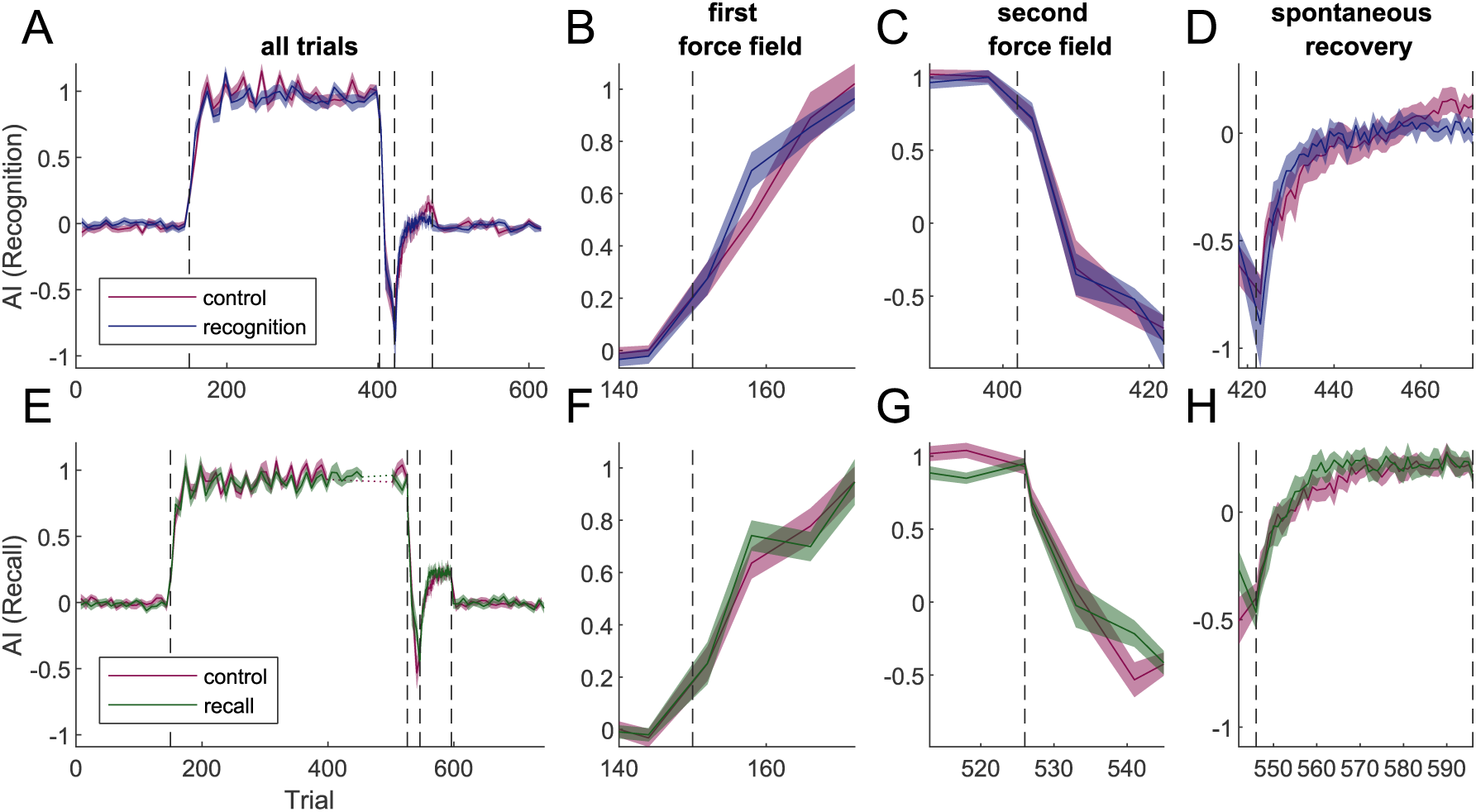
Adaptation index over trials for the memory (green and blue) and control (purple) tasks for the recognition group (**A-D**) and the recall group (**E-H**). **A, E**. AI for all trials. **B, F**. AI for trials around the first force field, **C, G**. AI for trials around the second force field. **D, H**. AI during trials in the spontaneous recovery phase.

We next adopted a modelling approach to interpret these subtle differences in spontaneous recovery, and to detect whether there was a potential impact that may have been hidden in the group means. To this end, we used a hierarchical Bayesian dual-rate model to simulate the AI of individual participants in the two groups (recognition and recall). The learning and retention parameters are drawn from population distributions with intervening recognition or recall tasks modelled as scaled versions of those for the control task (see Methods). We assumed that, at best, the declarative memory task does not affect the motor adaptation, meaning that scaling distributions were taken between 0 and 1.

Figure 5 shows the pooled parameter distributions for both groups (A: Recognition, B: Recall), and mode changes at the individual level are illustrated as connected dots (see Table 1 for 95% HDIs). While the distributions of the learning rates (second and fourth panels in A and B) seem to largely overlap between the declarative memory and the vowel counting task (control), the retention rates of both the fast and slow process show a mode shift in both the Recognition and Recall group. For the slow process, this mode shift is similar in the two groups. For the fast process, this shift appears larger in the recall than recognition group.

**Figure 5.**
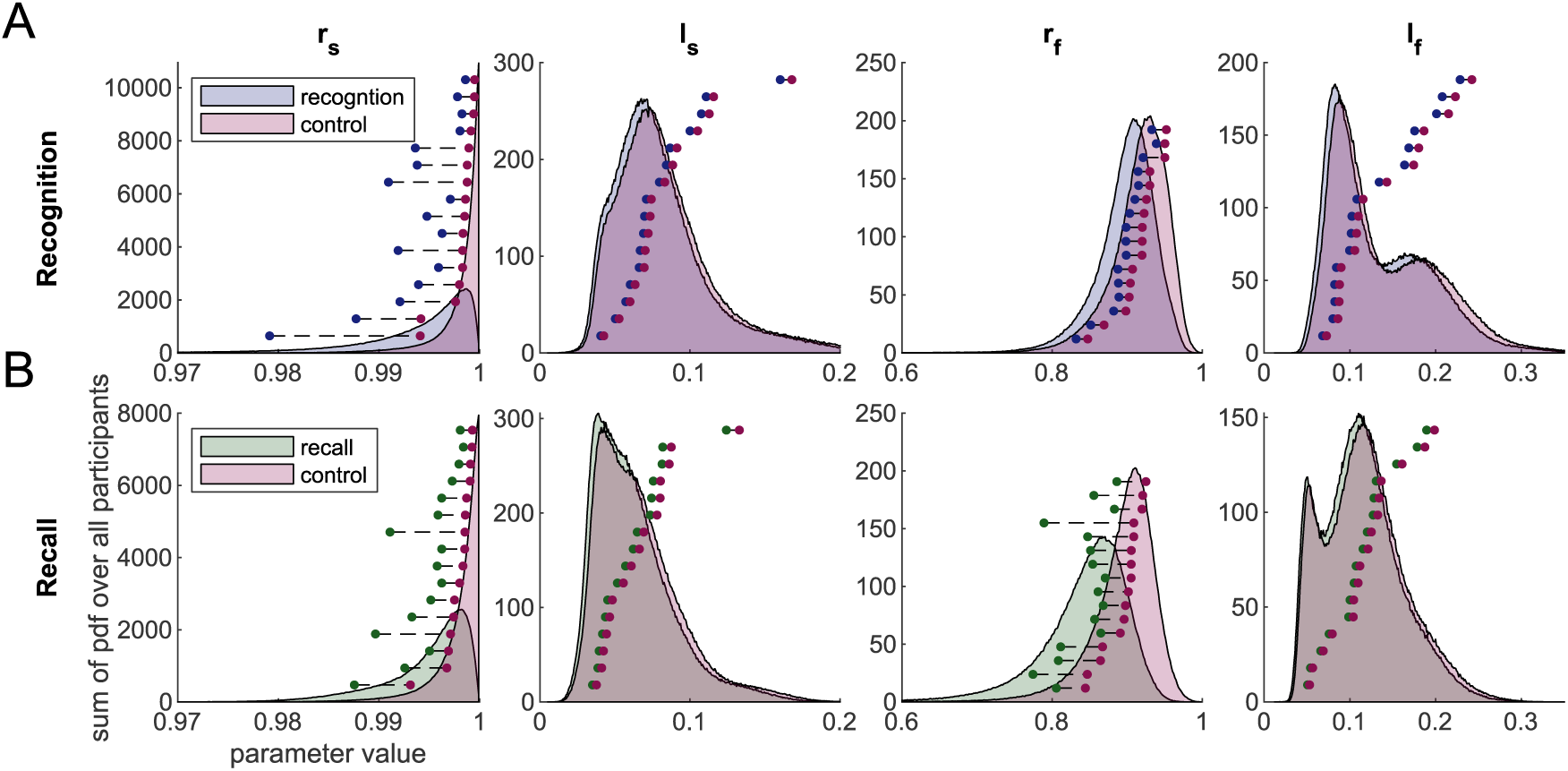
Learning (l) and retention (r) parameters of the slow (s) and fast process (f). **A.** Summarized parameter distributions (sum over individual participant distributions) for the recognition and control task. **B.** Summarized parameter distributions for the recall and control task. Dots represent the modes of the distributions of individual participants sorted by the control values.

To investigate whether these mode shifts are due to the intervening declarative memory task or whether they are a result of task or model structure, we next compared the scaling distributions inferred from the data to the simulated scaling distributions (see Methods). Figure 6 shows the group level scaling distributions for the four model parameters for the two experimental groups (A: Recognition; B: Recall; see Table 1 for 95% HDIs). The posterior distributions of the scaling parameters inferred from simulated data without scaling (control) are depicted in grey. For the slow and fast learning rate (second and fourth panels in A and B), as well as the slow retention rate (first panel), the posterior distributions of the data almost fully overlap with these simulated distributions, in both groups. Only the modes of the fast retention rates in the two groups, particularly in the Recall group, seem substantially pulled away from one, suggesting an impairment in motor adaptation due to the intervening declarative memory task.

**Figure 6.**
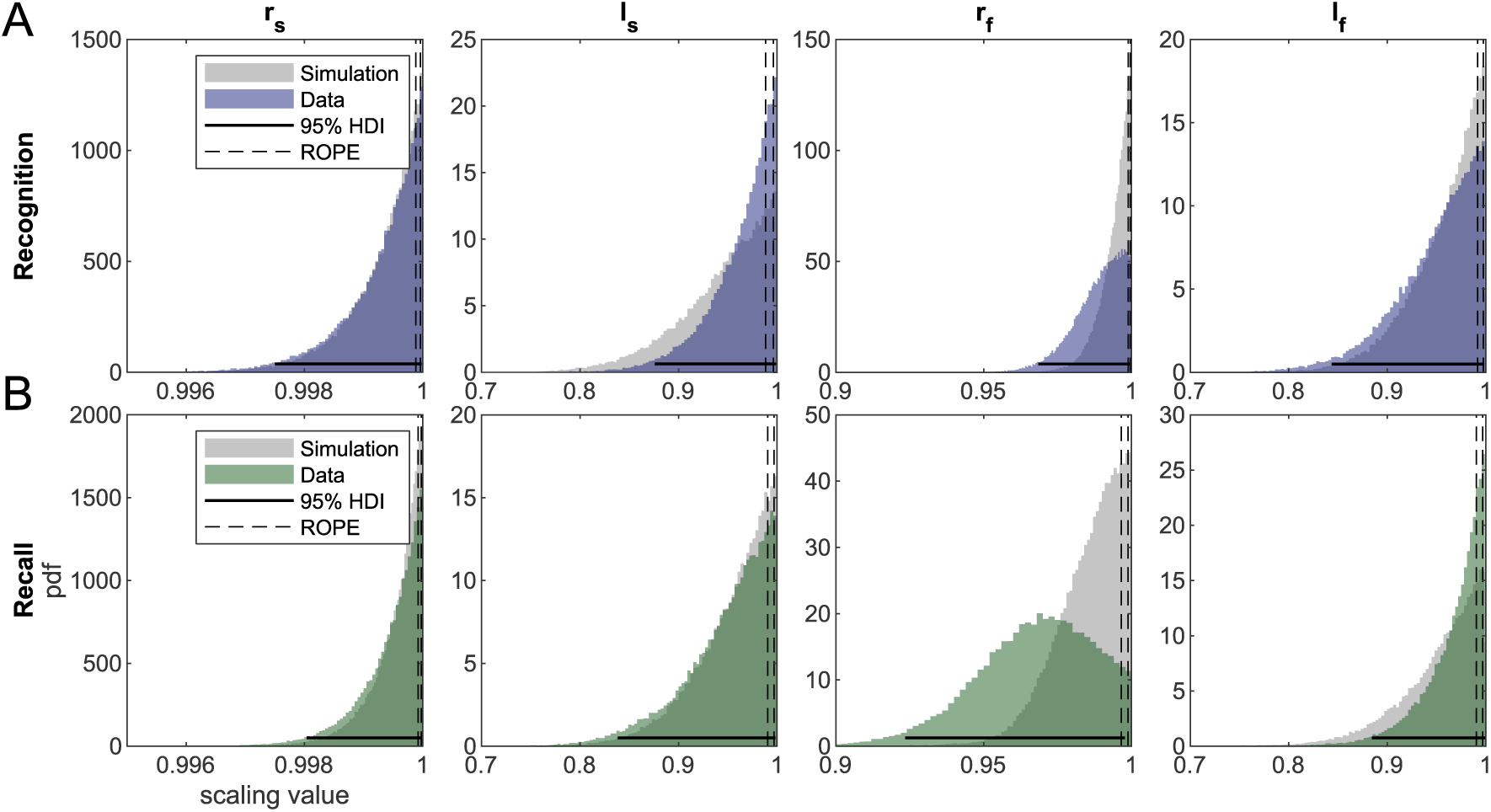
Posterior distribution of the scaling. for the hyperparameter of the learning and retention parameters based on the simulations (grey) and the data (**A.** Recognition group, blue; **B.** Recall group, green) Note that the ROPEs are represented by the two dashed lines, and the 95% HDI of the data is represented by the solid black horizontal line.

To test whether these shifts are of significance, we performed a ROPE analysis (see Methods). We compared the posterior distributions of the scaling of the learning and retention rates as obtained from the data to the posterior distributions of these hyperparameters obtained from the simulations (i.e., colored distribution and grey distributions in Figure 6). The scaling parameter for the learning and retention rates of the slow process and the learning rate of the fast process did not differ significantly in both groups since the 95% HDI (Figure 6, black line) of the posterior distribution of the respective data fully overlaps with the ROPE (Figure 6, dashed lines). According to the ROPE, the parameters are significantly equal. The scaling parameter of the fast retention rate showed a partial overlap of the 95% HDI with the ROPE, suggesting that we must remain undecided on whether this parameter differs significantly from one.

Finally, to assess how well simulated behavior based on parameter estimates mimics the participant data, we performed a posterior predictive check. We randomly drew 5000 iterations of the posterior distributions of the participants and simulated the model predictions for the control task. Figure 7 shows the posterior predictive AI (thin grey lines) as well as its expected mean (black line), the mean predicted fast (blue) and predicted slow (red) process, superimposed on data of all participants and tasks (pink ± SEM).

**Figure 7.**
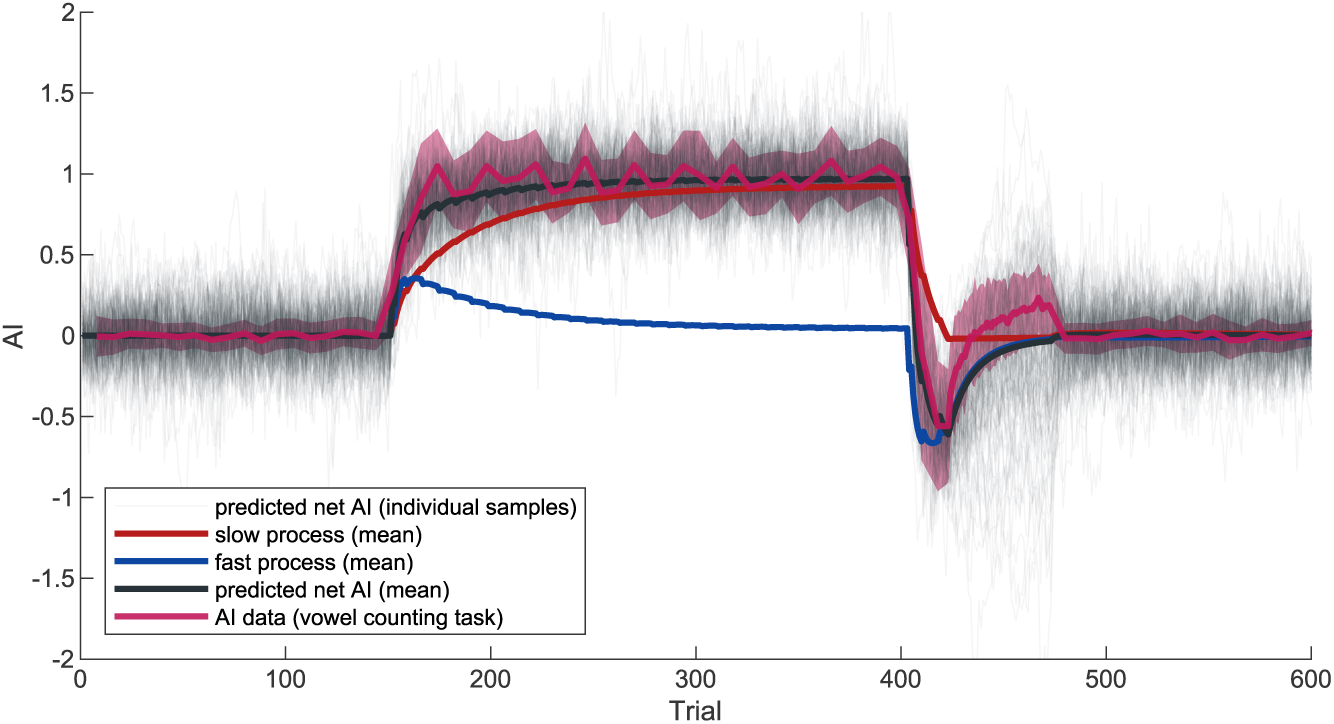
Posterior predictive check. Posterior predictive check on the control task and control data. Grey thin lines show 500 individual samples. The black line shows the average over all simulations (n = 5000). The average predicted state of the fast process is represented in blue, the average predicted state of the slow process is presented in red. The pink line with shaded area (±SEM) represents the data of the control task of all participants (n = 32).

## Discussion

We investigated the effect of a declarative memory task (word-list learning) on a well-established motor adaptation protocol that evokes spontaneous recovery. We assessed the trial-to-trial adaptation performance at the kinematic level, using the hand-path error, and at the dynamic level, using the adaptation index. Although the learning curves of both measures do not differ much compared to those collected with a control task (vowel counting), the AI suggests slight differences in the initial spontaneous recovery phase that is consistent across the two memory groups. Guided by a hierarchical Bayesian version of the dual-rate adaptation model, which advances the model proposed by Smith et al. (2006), we reverse engineered the behavioral outcome of each participant as the results of two adaptive process, one that responds quickly to error, but rapidly forgets and another that learns slowly from error but has strong retention. With this modelling approach, the estimate of the parameters of a single participant is based on group level hyperparameters describing these processes (Kruschke, 2015). By applying scaling distributions to the individual participants learning and retention rates from the control task, we could then infer to what extent the dual-rate parameters were impaired by the declarative memory task. While the fast retention rate visually appeared to be affected by the declarative memory task (Figure 5), applying the HDI+ROPE decision rule did not characterize this as conclusive evidence. For the other three parameters, this decision rule characterized them as significantly equal to those in the control task. In the following, we will discuss the implications of these findings.

Our modelling results do not provide strong evidence for interference of the declarative memory task on motor adaptation; if anything, there is a hint of a retrograde effect. This contrasts with previous work by Keisler and Shadmehr (2010), showing that a declarative memory task performed after a brief period of motor adaptation (for example, after the second force field in a spontaneous recovery paradigm), reduced the aftereffects of the briefly introduced perturbation in an error-clamp block. They explained their results by retrograde interference with the retention rate of the fast process. Their effects occurred in the initial phase of the error-clamp block, where our results also hinted at potential differences. It is important to point out that in their study the memory task was only carried out after the adaptation to the second force field, introducing retrograde interference before the spontaneous recovery phase, while in our paradigm the declarative memory tasks intervened continuously throughout the exposure of the two fields, potentially creating anterograde and retrograde interference throughout the adaptation and before the spontaneous recovery phase. Based on our results, it could be suggested that the fast process in motor adaptation is more susceptible to retrograde than anterograde interference effects. The subtle difference between the control and memory task in the initial phase of spontaneous recovery hints at similar interference characteristics as Keisler and Shadmehr (2010) found, but on a smaller scale. If motor adaptation is to share resources with the declarative memory system one would expect that the retrograde interference in our study would have a continuous and interactive effect on adaptation (see supplemental Figure S1), which was not clearly present in the behavioral data.

In this study, we used a declarative memory task – word-list learning – to interfere with the declarative component of motor adaptation. In other studies, the declarative or explicit component of motor adaption was directly measured (Schween et al., 2020; Taylor et al., 2014). For example, Schween et al. (2020) inferred the explicit component of force field adaptation by instructing participants to push (or not to push) against the force. Their results hint at a benefit from explicit strategy use in force field adaptation. Taylor et al. (2014) investigated whether visuomotor adaptation of reaching movements included an explicit component by asking participants for their aiming direction. When subtracting the aiming direction from the actual movement angle of the hand one can infer the implicit component. It was found that the implicit and explicit component mimic the dynamics of the fast and slow processes, respectively (McDougle et al., 2015). In line with their results, our results show no impact of the intervening word learning task on the slow process. However, our results yield no clear significant impact (only a hint) of the declarative memory task on the fast process of motor adaptation either.

Hadjiosif and Krakauer (2021) suggested that probing the explicit strategy during adaptation could increase the use of an explicit strategy, which otherwise may stay unused. In line with this, the introduction of explicit strategy use alters the pattern of motor adaptation, including faster initial learning (Benson et al., 2011; Maresch et al., 2021; Schween et al., 2020). In our study, the declarative memory task was of a different nature than the motor adaptation task itself and could therefore be assumed to not yield any motor benefits but instead only have adverse interference effects. Clearly, the interference of our declarative memory task was only subtle, and only retrogradely, but we cannot claim statistical significance. It could be argued that the explicit memory system is only used as a back-up system when implicit learning falls short. For example, De Brouwer et al., (2018) made use of the gaze direction as a more indirect way to quantify the explicit strategy in motor adaptation. They show that gaze direction served that function better in participants who also had to indicate the explicit aim verbally compared to those without such verbalization. Their results suggest that an explicit strategy use in motor adaptation does not occur naturally, at least not in every individual. In this vein, it is possible that only a few individuals used an explicit strategy in our paradigm, which could explain why significance was not reached on a group level.

Other evidence for an explicit memory component in motor adaptation comes from studying participants with decreased declarative memory capacity. For example, McDougle et al. (2022) reported an increase in the fast retention rate in a single patient with MTL damage. On the other hand, Sutter et al. (2024) shows a decrease in the fast retention rate in group of patients with Alzheimer’s dementia. Furthermore, Trewartha et al. (2014) showed that poor explicit memory in elderly participants was associated with reduced retention by the fast process. All these studies have in common that they tested adaptation in an older and/or clinical population. It could be argued that with age we rely more on explicit strategies to solve motor adaptation tasks (Oosterman et al., 2021). To test this hypothesis, future studies should focus on a geriatric population, using our paradigm to see whether the declarative memory task has a stronger impact in this population.

An alternative explanation of our results is that our memory manipulation did not sufficiently tax the declarative memory system to interfere with motor adaptation. Sing and Smith (2010) showed that anterograde interference increases with exposure to the task (i.e., the number of trials). One could suggest that the 5 cycles of the same word recall task was not sufficient to impair motor adaptation in our paradigm, even though it was not undemanding as participant’s recall performance did improve from cycle to cycle.

In this study, we conjecture impaired learning to anterograde interference and impaired retention to retrograde interference. Recently, Hamel et al. (2022) showed that anterograde interference depends on the overlap in neural correlates involved in the interfering tasks. If two tasks share no neural structures, their level of interference is low, while in case of overlapping neural correlates, interference is more substantial. In an elementary view, motor adaptation is primarily related to the cerebellum (Lang & Bastian, 1999; Statton et al., 2018) while the declarative memory system largely relies on the MTL. Therefore, the lack of anterograde interference in our study is in line with the idea that interference only occurs if there is overlap in the corresponding neural structures. Although not tested in Hamel et al. (2022), if their hypothesis extends to retrograde interference as well, it could explain the non-significant impairment of the fast retention rate.

Recently, Heald et al. (2021, 2023) introduced the notion of contextual interference to explain adaptation behavior. Following their model, the behavioral pattern of spontaneous recovery occurs due to the expression of two contextual memories: one corresponding with the first learned force field with high probability of re-occurring, and one corresponding to the briefly introduced second force field with lower probability of reoccurring. Both memories are represented by a single state variable, in contrast to the two states in the dual-rate model. When error feedback is removed, the probability of being in the initial force field is higher than being in the second. Within this notion, it could be argued that the declarative memory task did not directly interfere with motor adaptation but rather with the process of inferring the current context. In comparison, the application of a word-learning task after a period of sole motor learning by Keisler and Shadmehr (2011) could have caused a new memory context, which decreased the probability of the other two contexts from re-occurring. Since the second force field was only introduced briefly, the probability that context would reoccur was impacted more strongly than the initially learned high probability context. In line with this idea, Ning and Wright, (2023) showed that anterograde interference is larger in a blocked design than in a continuous design in perceptual learning. In our study, the declarative memory context and the motor adaptation context occurred in parallel and therefore showed less interference on the expression of the first and second force field.

## Grants

This work was supported by an internal grant from the Donders Centre for Cognition. W.P.M. is additionally supported by the following grants: NWA-ORC-1292.19.298, NWO-SGW-406.21.GO.009 and Interreg NWE-RE:HOME.

## Author contributions

J.L.R., L.P.J.S. and W.P.M. conceived and designed research; J.L.R. performed experiments; J.L.R. analyzed data; J.L.R., L.P.J.S. and W.P.M. interpreted results of experiments; J.L.R. prepared figures; J.L.R. drafted manuscript; J.L.R., L.P.J.S. and W.P.M. edited and revised manuscript; J.L.R., L.P.J.S. and W.P.M. approved final version of manuscript.

## Data availability statement

Upon publication, all data and code will be made publicly available on the Radboud Data Repository.

## Supplemental figure S1

**Figure S1.**
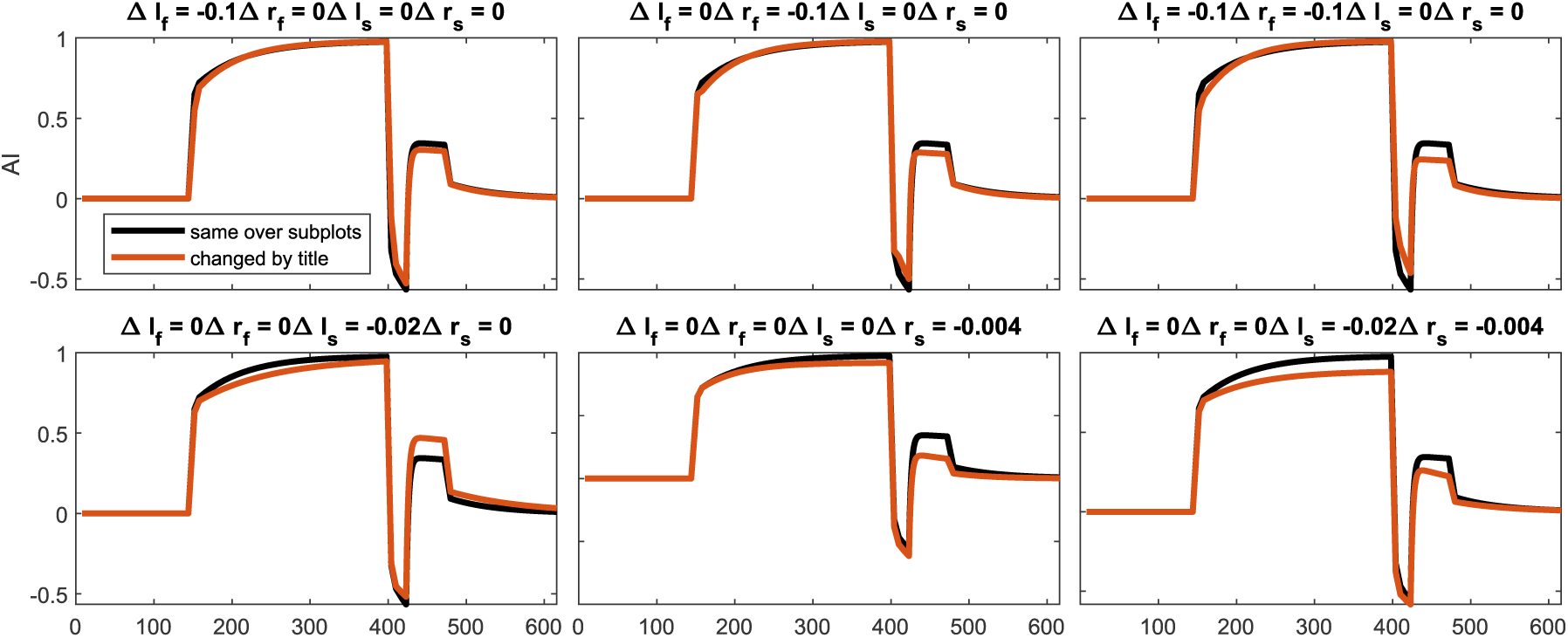
Effects of impaired parameters on net adaptation. Model predictions during a spontaneous recovery paradigm. Same parameter values were used for the predictions color-coded in black. Top row shows an impaired fast process, bottom row an impaired slow process. First column shows impaired learning, second column impaired retention, and the third column the combination of impaired learning and retention. The black line shows adaptation behavior of a learner with a fast learning rate of 0.6, a fast retention rate of 0.7, a slow learning rate of 0.05, and a slow retention rate of 0.995. The orange line represents the adaptation behavior with impairment.

